# The complete chloroplast genome sequence of *Akebia trifoliata* and a comparative analysis within the Ranales clade

**DOI:** 10.1101/2023.01.18.524534

**Authors:** Ruixian Wang, Qu Chu, Gang Meng, Chuan Li

**Affiliations:** School of Modern Agriculture and Biotechnology, Ankang University, No. 92 Yucai Road, Hanbin District, Ankang, 725000, Shaanxi Province, China; Shaanxi Key Laboratory of Sericulture; Ankang University, No. 92 Yucai Road, Hanbin District, Ankang, 725000, Shaanxi Province, China; Institute of Shaanxi Selenium-enriched Recycling Agricultural Development, No. 92 Yucai Road, Hanbin District, Ankang, 725000, Shaanxi Province, China

**Author notes:** Corresponding Author: Chuan Li.

## Abstract

The complete nucleotide sequence of the *Akebia trifoliata* chloroplast (cp) genome was reported and characterized in this study. The cp genome is a closed circular molecule of 157 949 bp, composed of a pair of IR regions of 26 149 bp, one LSC region of 86 595 bp, and one SSC region of 19 056 bp. The GC contents of the LSC, SSC, and IR regions, and the whole cp genome are 37.11%, 33.62%, 43.08% and 38.66%, respectively. The cp genome contains 130 predicted functional genes, including 85

PCGs of 79 107 bp, 37 tRNA and 8 rRNA genes of 11 851 bp. 168 SSRs and 23 long repeats were identified in the cp genome. The results revealed that 21 891 codons characterize the coding capacity of 85 protein-coding genes in *Akebia trifoliata*, 10.84% and 1.20% of the codons coded for leucine and cysteine respectively. The usage of the start codon exhibited no bias in the *A. trifoliata* cp genome. Phylogenetic analysis suggest that *A. trifoliata* is most closely related to *S. japonica*, which then formed a cluster with *N. dormestica* and *M. saniculifolia* to form subgroup of Ranunculales. Our study provides information on mangrove plant species in coastal intertidal zones.

## Introduction

*Akebia trifoliata* is a wild perennial woody liana, and monoecious with flowers functionally unisexual and self-incompatible [1, 2]. It belongs to the Lardizabalaceae family, that is mainly distributed in the eastern part of Asia, and is also commonly known in China as August melon, Bayuezha, and wild banana [3]. Its fresh fruit, commonly called ‘Ba-Yue-Gua’ in China, has long been consumed by the local people as a delicious food [4]. *A. trifoliata* has been used in traditional Chinese medicine (TCM) for more than 2000 years [5]. The air-dried stems and fruits of *A. trifoliata* have traditionally been used in China for centuries as an antiphlogistic, antineoplastic and diuretic agent [6]. To date, phytochemical studies have revealed structurally diverse of triterpenes, triterpene saponins, phenolics and lignans from this plant, and some of them displayed significant biological activities [7-11]. Owing to its economic and medicinal values, *A. trifoliata* is currently being widely cultivated and rapidly developed as a commercial species in many regions of China.

Chloroplast serves as the metabolic center of plant life by converting solar energy to carbohydrates through photosynthesis and oxygen release [12]. The chloroplast genome consists of 120-130 genes that divided into three functional categories, protein-coding genes, introns and intergenic spacers. They comprise a single circular chromosome, typically ranging in size from 107 kb to 218 kb [13, 14]. Chloroplast genomes have a quadripartite structure, with a pair of inverted repeats (IRs) separated by one large and one small single copy region [15]. Several plant chloroplast genomes also show significant structural rearrangements, with evidence of the loss of IR regions or entire gene families [16, 17]. Additionally, the presence of IRs might stabilize the chloroplast genomes organization [18, 19]. Approximately 800 complete chloroplast genomes from a variety of land plants have been retained in the National Center for Biotechnology Information (NCBI) organelle genome database since the first chloroplast genome of tobacco [20] and liverwort [21], which were sequenced simultaneously in 1986. The sequenced chloroplast genomes have improved our understanding of plant biology and evolutionary relationships.

In this study, the complete chloroplast genome of *A. trifoliata* was sequenced and analyzed based on illumina high-throughput sequencing technology. In addition to describing the plastic features of the chloroplast genome, the gene content, repeat structures and sequence divergence with other reported species in the Ranunculales order were compared. We also presented results of a phylogenetic analysis of protein sequences from *A. trifoliata* and 40 other plant species. The complete chloroplast genome of *A. trifoliata* will improve our understanding of the evolution relationships of genera in the Ranunculales order.

## Materials and Methods

### Sampling and DNA sequencing

Based on its morphological characteristics, the *A. trifoliata* species, in its natural habitat of the **Tsinling Mountains (106°55′19″E, 34°14′29″N)**, was confirmed by Professor Chuan Li and Yunwu Peng (Ankang University). The plant was then transplanted and preserved in the School of Modern Agriculture and Biological Science and Technology, Ankang University. Approximately, 20g fresh leaves were sampled from a single *A. trifoliata* plant, and the cpDNA was extracted using a modified high salt method. Approximately 5-10 μg of cpDNA was sheared, followed by adapter ligation and library amplification. Then, the fragmented cpDNAs were sequenced using the Illumina Hiseq 2000 platform. The obtained sequences were assembled using SOAP de novo software and reference-based approaches in parallel. The obtained cp genome regions with ambiguous alignments were manually trimmed and considered as gaps. Gaps were filled using the read over-hangs at margins and the PCR method. The process was repeated fold, and the minimum fold of the final assembled *A. trifoliata* cpDNA reached approximately 1126-fold.

### Genome annotation

The tRNA, rRNA, and protein-coding genes (PCGs) in the assembled genome were predicted and annotated using Dual Organellar GenoMe Annotator with default parameters. The locations of tRNAs were then confirmed using tRNAscan-SE software, version 1.21, specifying mito/cpDNA as the source. The rRNA genes were verified using BLASTN searches against the database of published cp genomes. The positions of the start and stop codons, or intron junctions of PCGs, were verified using the BLASTN searches and sequin program with Plastid genetic code. The genome map was drawn by OGDraw v1.2.

### Simple sequence repeat (SSR) and long repeats analysis

Distributed throughout the genome, SSRs are repeat sequences with a typical length of 1-6 bp that are generally considered to have a higher mutation rate than neutral DNA regions. The distributions of SSRs in the chloroplast genome were predicted by using the microsatellite search tool MISA [22] with the following parameters: ≥10 for mononucleotide repeats, ≥5 for dinucleotide repeats, ≥4 for trinucleotide repeats, and ≥3 for tetranucleotide repeats, pentanucleotide repeats, and hexanucleotide repeats. For analysis of repeat structures, REPuter [23] was used to visualize forward, reverse, palindrome and complement sequences of size ≥30 bp and identity ≥90% in the cp genome. All repeats recognized were manually verified, and redundant results removed.

### Comparative Genome analysis

To investigate the sequence divergence of the chloroplast genome among the analyzed Ranunculales species, the whole chloroplast genome sequences of *Nandina domestica* of Berberidacae, *Euptelaea pleiosperma* of Eupteleaceae, *Stephania japonica* of Menispermaceae, *Megaleranthis saniculifolia* of Ranunculaceae and *Papaver somniferum* of Papaveraceae were analyzed using the mVISTA program in the Shuffle-

LAGAN mode [24], and the *Euptelea pleiosperma* annotations were used as references. The differences in the chloroplast genome length, LSC length, SSC length, GC content, encoding gene types and gene numbers among these 6 species were analyzed. The LSC/IR/SSC boundaries among the species were determined by comparative analysis to explore the variation in these angiosperm chloroplast genomes.

### Phylogenetic analysis

To illustrate the phylogenetic relationship of the *A. trifoliata* with other major Ranales clades, 40 complete cp genomes were downloaded from GenBank (Table S1). *Sapindus mukorossi* and *Acer buergerianum* from Sapindales were used as outgroups. Then, 72 PCGs found in all of the species were extracted from the selected cp genomes. The amino acid sequences of each of the PCGs were aligned using MSWAT (http://mswat.ccbb.utexas.edu/) with default settings, and back translated to nucleotide sequences. Phylogenetic analyses were performed using the concatenated nucleotide sequences and RAxML 7.2.6 software by the maximum likelihood (ML) method. RAxML searches relied on the General Time Reversible model of nucleotide substitution with the gamma rate model (GTRGAMMA). A bootstrap analysis was performed with 1000 replications using non-parametric bootstrapping as implemented in the “fast bootstrap” algorithm. The absent genes were finally mapped on to the ML phylogenetic tree.

## Results

### The overall structure and general features of the *A. trifoliata* cp genome

The cp genome of *A. trifoliata* is a closed circular molecule of 157 949 bp (GenBank accession number: ON021949), composed of a pair of IR regions (IRa and IRb) of 26 149 bp, one LSC region of 86 595 bp, and one SSC region of 19 056 bp. It has an overall typical quadripartite structure that resembles the majority of land plant cp genomes. The GC contents of the LSC, SSC, and IR regions, and the whole cp genome are 37.11, 33.62, 43.08 and 38.66 %, respectively (Table 1). The cp genome encodes 130 predicted functional genes, including 85 PCGs of 79 107 bp, 37 tRNA and 8 rRNA genes of 11 851 bp. Among the 130 genes, 114 are unique genes and 19 are duplicated genes in the IR regions. 16 genes have one intron (10 PCGs and 6 tRNA genes) and 2 PCGs have two introns (*clpP* and *ycf3*). Like most other land plants, a maturase K gene (*matK*) is located within the intron of *trnK, rps12* is trans-spliced, with its two 3′ end residues separated by an intron in the IR region, and the 5′ end exon is in the LSC region (Figure 1). The 8 rRNA genes were composed of two identical copies of 16S-23S-4.5S-5S rRNA gene clusters in the IR region. Each cluster was interrupted by two tRNA genes, *trnI* and *trnA*, in the 16S-23S spacer region.

**Table 1.** Base composition in the *A. trifoliate* chloroplast genome.

| Terms | A (%) | T (%) | C (%) | G (%) | length (bp) |
| --- | --- | --- | --- | --- | --- |
| LSC | 30.82 | 32.07 | 19.00 | 18.11 | 86595 |
| SSC | 33.18 | 33.20 | 17.61 | 16.01 | 19056 |
| IRa | 28.22 | 28.70 | 22.23 | 20.85 | 26149 |
| IRb | 28.70 | 28.22 | 20.85 | 22.23 | 26149 |
| Total | 30.32 | 31.01 | 19.67 | 18.99 | 157949 |
| CDS | 30.30 | 30.86 | 18.22 | 20.61 | 79107 |

**Figure 1.**
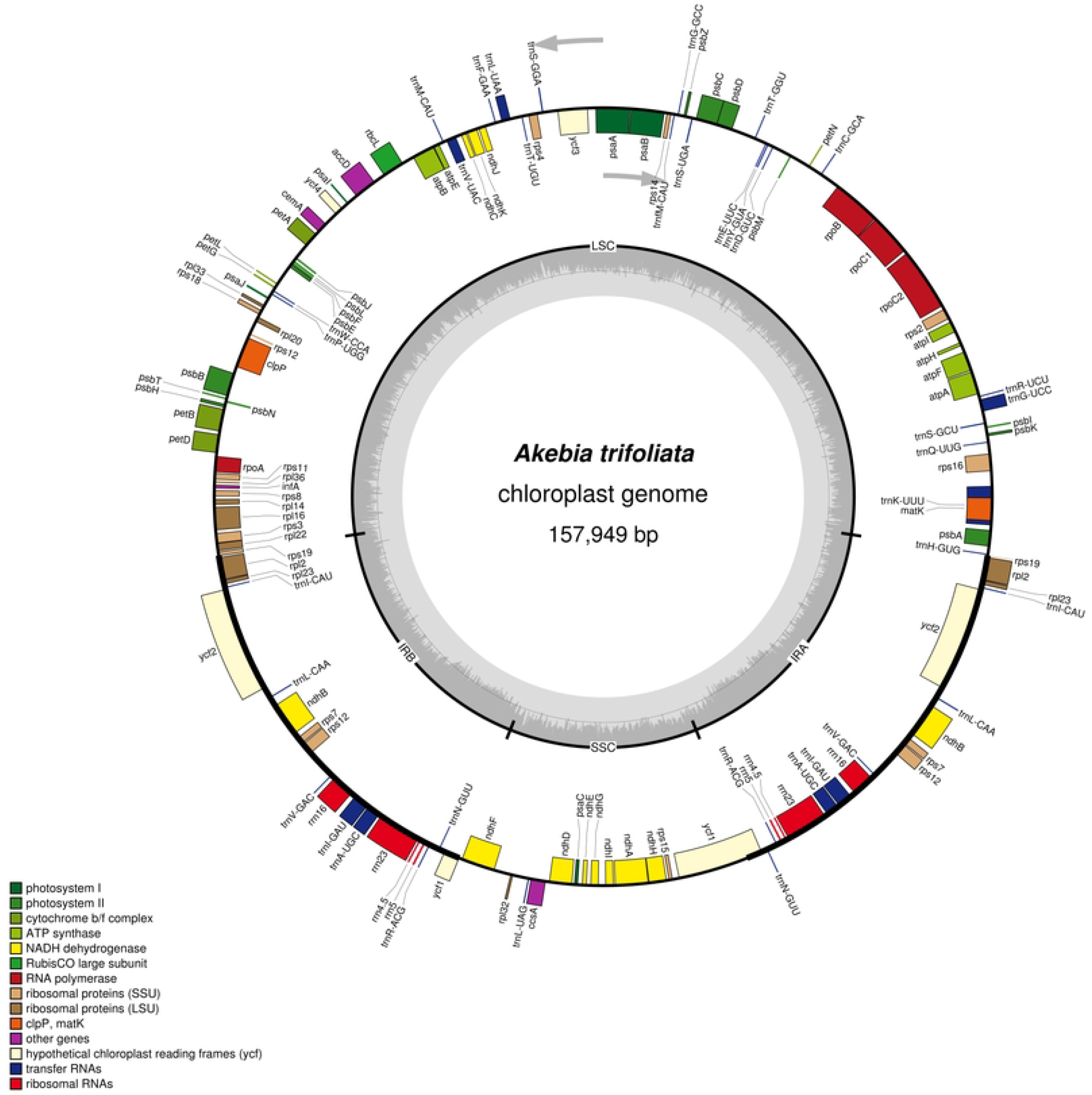
Gene map of the *A. trifoliata*. Genes lying outside of the outer layer circle are transcribed in the counterclockwise direction, whereas genes inside are transcribed in the clockwise direction. The colored bars indicate known different functional groups. Area dashed darker gray in the inner circle denotes GC content while the lighter gray shows to AT content of the genome. LSC large single-copy, SSC small-single-copy, IR inverted repeat.

### Codon Usage

Codon usage plays an important role in shaping chloroplast genome evolution. Based on the sequences of protein-coding genes (CDS), the codon usage frequency was estimated (Table 2). The result revealed that 21 891 codons characterize the coding capacity of 85 protein-coding genes in *A. trifoliata*. Of these codons, 2 373 (10.84%) were code for leucine and 263 (1.20%) for cysteine, which represented the maximum and minimum prevalent number of amino acids in the *A. trifoliata* chloroplast genome, respectively. A- and U-ending codons were ordinary. The other amino acid codons in the cp genome preferentially end with A or U (RSCU > 1). This codon usage pattern is similar to those reported cp genomes. In addition, codons ending in A and/or U accounted for 71.96% of all CDS codons. The majority of protein-coding genes in land-plant cp genomes employ standard ATG initiator codons. The use of the start codon exhibited no bias (RSCU = 1) in the *A. trifoliata* cp genome.

**Table 2.** Codon usage of *A. trifoliate* chloroplast genome.

| Amino acids | Codon | No. | RSCU | Amino acids | Codon | No. | RSCU |
| --- | --- | --- | --- | --- | --- | --- | --- |
| Phe | UUU | 751 | 1.22 | Ala | GCC | 191 | 0.64 |
| Phe | UUC | 486 | 0.75 | Ala | GCA | 362 | 1.05 |
| Leu | UUA | 689 | 1.76 | Ala | GCG | 149 | 0.41 |
| Leu | UUG | 500 | 1.22 | Tyr | TAT | 664 | 1.46 |
| Leu | CUU | 499 | 1.26 | Tyr | TAC | 167 | 0.39 |
| Leu | CUC | 172 | 0.45 | His | CAT | 434 | 1.19 |
| Leu | CUA | 325 | 0.77 | His | CAC | 145 | 0.40 |
| Leu | CUG | 188 | 0.55 | Gln | CAA | 607 | 1.45 |
| Ile | AUU | 939 | 1.53 | Gln | CAG | 205 | 0.37 |
| Ile | AUC | 418 | 0.60 | Asn | AAT | 817 | 1.32 |
| Ile | AUA | 596 | 0.87 | Asn | AAC | 244 | 0.46 |
| Val | GUU | 475 | 1.44 | Lys | AAA | 821 | 1.36 |
| Val | GUC | 154 | 0.48 | Lys | AAG | 304 | 0.41 |
| Val | GUA | 474 | 1.44 | Asp | GAT | 732 | 1.41 |
| Val | GUG | 197 | 0.58 | Asp | GAC | 207 | 0.41 |
| Ser | UCU | 457 | 1.54 | Glu | GAA | 878 | 1.35 |
| Ser | UCC | 295 | 0.99 | Glu | GAG | 313 | 0.50 |
| Ser | UCA | 366 | 1.32 | Cys | TGT | 188 | 1.13 |
| Ser | UCG | 175 | 0.47 | Cys | TGC | 75 | 0.27 |
| Ser | AGU | 344 | 1.28 | Arg | CGT | 324 | 1.46 |
| Ser | AGC | 108 | 0.32 | Arg | CGC | 88 | 0.33 |
| Pro | CCU | 371 | 1.44 | Arg | CGA | 320 | 1.46 |
| Pro | CCC | 190 | 0.70 | Arg | CGG | 99 | 0.37 |
| Pro | CCA | 285 | 1.22 | Arg | AGA | 418 | 1.60 |
| Pro | CCG | 112 | 0.52 | Arg | AGG | 161 | 0.55 |
| Thr | ACU | 460 | 1.60 | Gly | GGT | 555 | 1.32 |
| Thr | ACC | 237 | 0.72 | Gly | GGC | 169 | 0.39 |
| Thr | ACA | 373 | 1.22 | Gly | GGA | 642 | 1.57 |
| Thr | ACG | 134 | 0.41 | Gly | GGG | 256 | 0.62 |
| Ala | GCU | 586 | 1.80 |  |  |  |  |

### Analyses of simple sequence repeats (SSRs) and long repeats

A total of 168 SSR loci, harboring 220 bp in length, were detected in the *A. trifoliata* cp genome. 33 of them present in compound formation, and the number of mono-, di-, tri- and tetra-nucleotide repeats were 126, 35, 4 and 3 respectively. 120 of the mononucleotides, 16 dinucleotides, all of the trinucleotides and 1 trinucleotide were composed of A and T nucleotides, with a higher AT content (87.73%) in these sequences than in the cp genome. Our findings agreed with the observation that chloroplast SSRs were generally composed of polyadenine (poly A) and polythymine (poly T), and rarely contained tandem guanine (G) and cytosine (C) repeats. Among the SSRs, 43 were located in IGS regions and 10 were found in coding genes, including *atpF, cemA, ndhF, rpoC2, atpB, rpoB*, and *ycf1*.

Repeat analysis revealed 23 long repeats, include 8 forward (direct) and 15 palindrome (inverted) (Table 3). More than half of the repeats were located in the intergenic or intronic regions. The majority of these repeats were between 30 and 73 bp. Short dispersed repeats are considered to be a major factor promoting cp genome rearrangements, which may facilitate intermolecular recombination and create diversity among the cp genomes in a population. Hence, the repeats identified in this study will provide valuable information to support investigation of the phylogeny of *A. trifoliate* population studies.

**Table 3.**
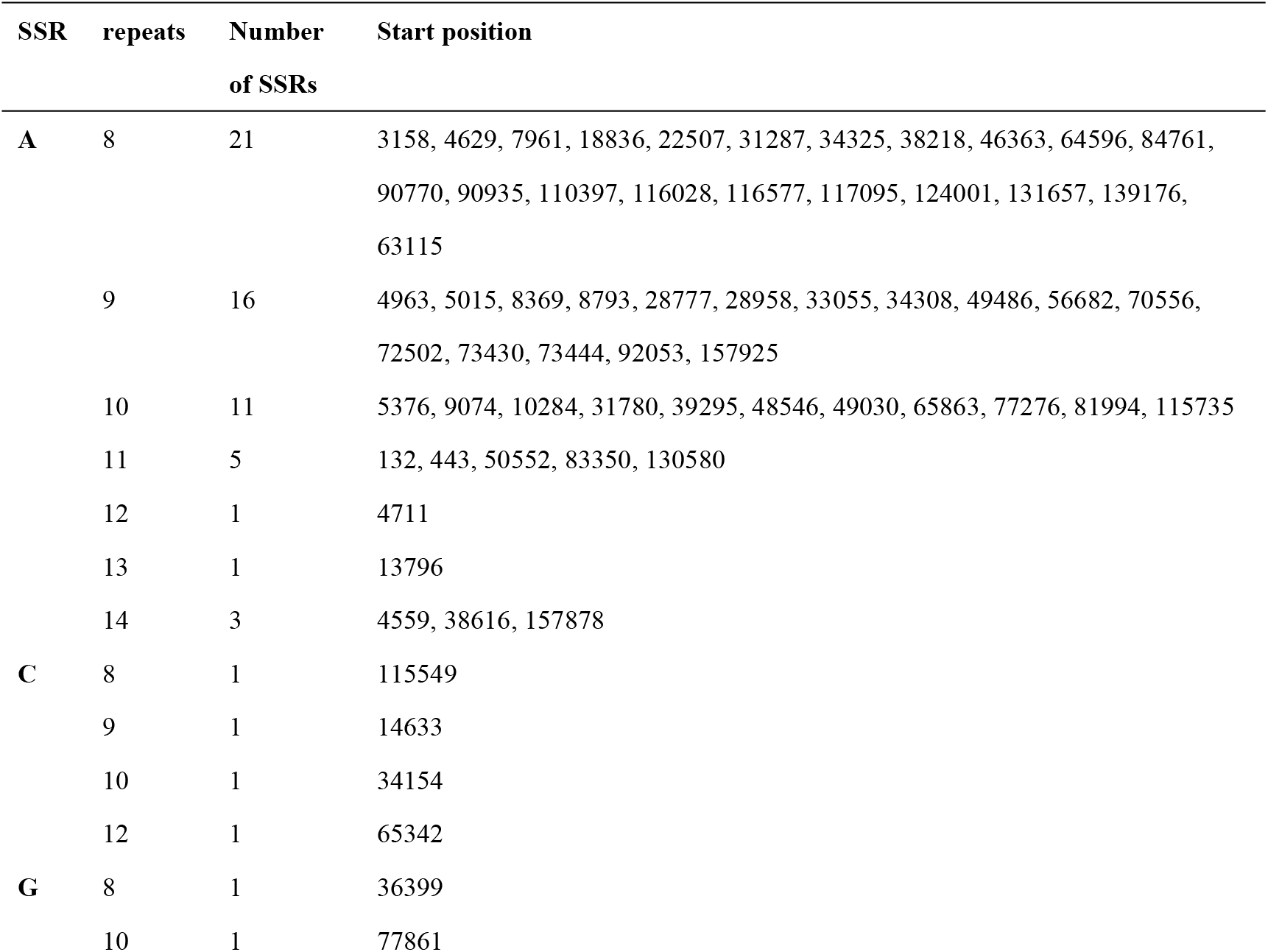
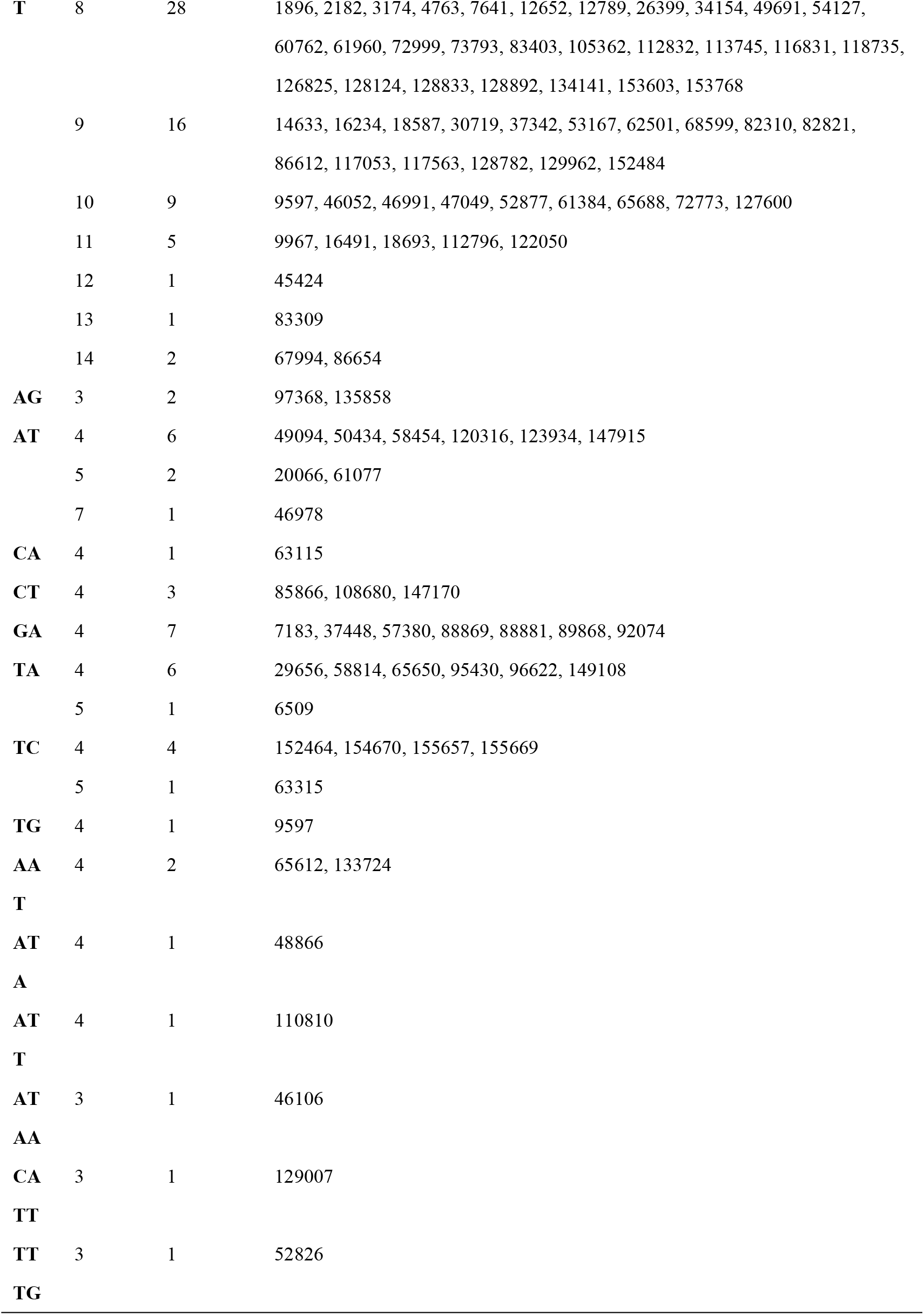
SSRs in *A. trifoliata* cp genome.

### IR contraction and expansion

Contraction and expansion at the borders of IR regions are common evolutionary events considered the main reason for size differences among cp genomes. For members of ranunculales species, we conducted an exhaustive comparison of four junctions, LSC-IRA (JLA), LSC-IRB (JLB), SSC-IRA (JSA), and SSC-IRB (JSB), between the two IRs (IRA and IRB) and the two single-copy regions (LSC and SSC). The results shown that that the studied locations are generally similar to those of all previously reported chloroplast genomes (Figure 2).

**Figure 2.**
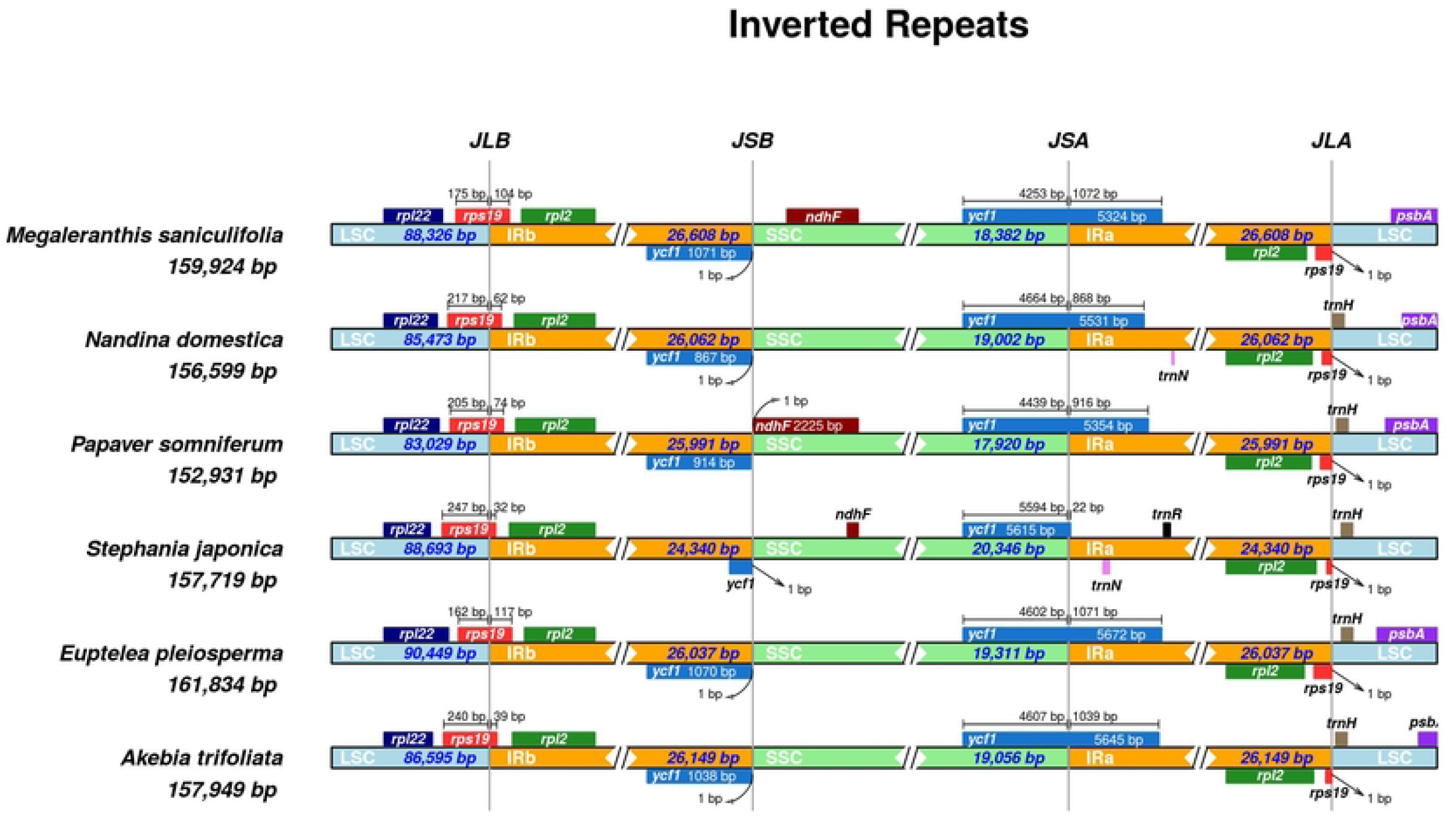
Comparisons of LSC, SSC, and IR region borders among six Ranunculales chloroplast genomes. The arrows indicated the location of the distance. This figure is not to scale.

Characteristically, there are four junctions in the chloroplast genomes of angiosperms, due to the presence of two identical copies of the inverted repeats. All chloroplast genomes appeared to be structurally similar with a typical quadripartite structure of two IRs separated by an LSC and an SSC. The whole genome sizes ranged from 152 931 (*P. somniferum*) to 161 834 (*E. plelosperma*).

There are two copies *rps19* genes in cp genomes. The JLB junction was placed in the rps19 region in all the cp genomes of Ranunculales species and outspread to different lengths (*M. sanlculifolia*, 175 bp; *N. domestica*, 217 bp; *P. somniferum*, 205 bp; *S. japonica*, 247 bp; *E. pleiosperma*, 162 bp; *A. trifoliata*, 240 bp) within the IRB region of all the genomes. The IRB region contained 104, 62, 74, 32, 117 and 39 bp of the *rps19* gene, respectively. Another rps19 gene near by the end of IRA region, and cut off by the JLA boundary.

For almost all the analyzed genomes, the *trnH* gene is the first gene in the LSC region with variable distance from JLA boundary, while *M. saniculifolia* has lost the trnH gene. Comparisons revealed that the *ndhF* gene have been lost in the *A. trifoliata, N. domestica* and *E. pleiosperma* cp genome.

### Comparative analysis of the chloroplast genomes among Ranunculales and phylogenetic analysis

The cp genome comparative analysis of several ranunculales species were conducted by the mVISTA program, with *Euptelea pleiosperma* as the reference sequence (Figure 3). Considerable similarities in genome composition and size were identified among the species. In these species, the *Papaver somniferum* has the smallest size of cp genome, and its size of LSC and IR region also the shortest (Table 4). The results of the comparison shown that the IR (A/B) regions exhibited fewer differences than the LSC and SSC regions. Moreover, the non-coding regions showed more variability than the coding regions (CNS), and the marked differences in regions among the six chloroplast genomes were evident in the intergenic spacers.

**Table 4.** Basic characteristics of the complete chloroplast genome of the six species. The GenBank No. of *N. domestica, E. pleiosperma, S. japonica, M. saniculifolia* and *P. somniferum* is NC_008336, KU204900, KU204903, NC_012615 and KU204905 respectively. All lengths are in bp.

| Species | Taxon | Genome size | LSC size | SSC size | IR size | Number of genes | Protein coding genes | tRNA | rRNA | GC (%) |
| --- | --- | --- | --- | --- | --- | --- | --- | --- | --- | --- |
| *N. domestica* | Berberidaceae | 156 599 | 85 473 | 19 002 | 26 062 | 134 | 79 | 30 | 8 | 38.3 |
| *E. pleiosperma* | Eupteleaceae | 161 834 | 90 449 | 19 331 | 26 037 | 132 | 79 | 30 | 8 | 38.6 |
| *S. japonica* | Menispermaceae | 157 717 | 88 693 | 20 346 | 24 340 | 132 | 79 | 30 | 8 | 38.2 |
| *M. saniculifolia* | Ranunculaceae | 159 924 | 88 326 | 18 382 | 26 608 | 131 | 79 | 30 | 8 | 38.0 |
| *P. somniferum* | Papaveraceae | 152 931 | 83 029 | 17 920 | 25 991 | 132 | 79 | 30 | 8 | 38.9 |
| *A. trifoliata* | Lardizabalaceae | 157 949 | 86 595 | 19 056 | 26 149 | 130 | 85 | 37 | 8 | 38.7 |

**Figure 3.**
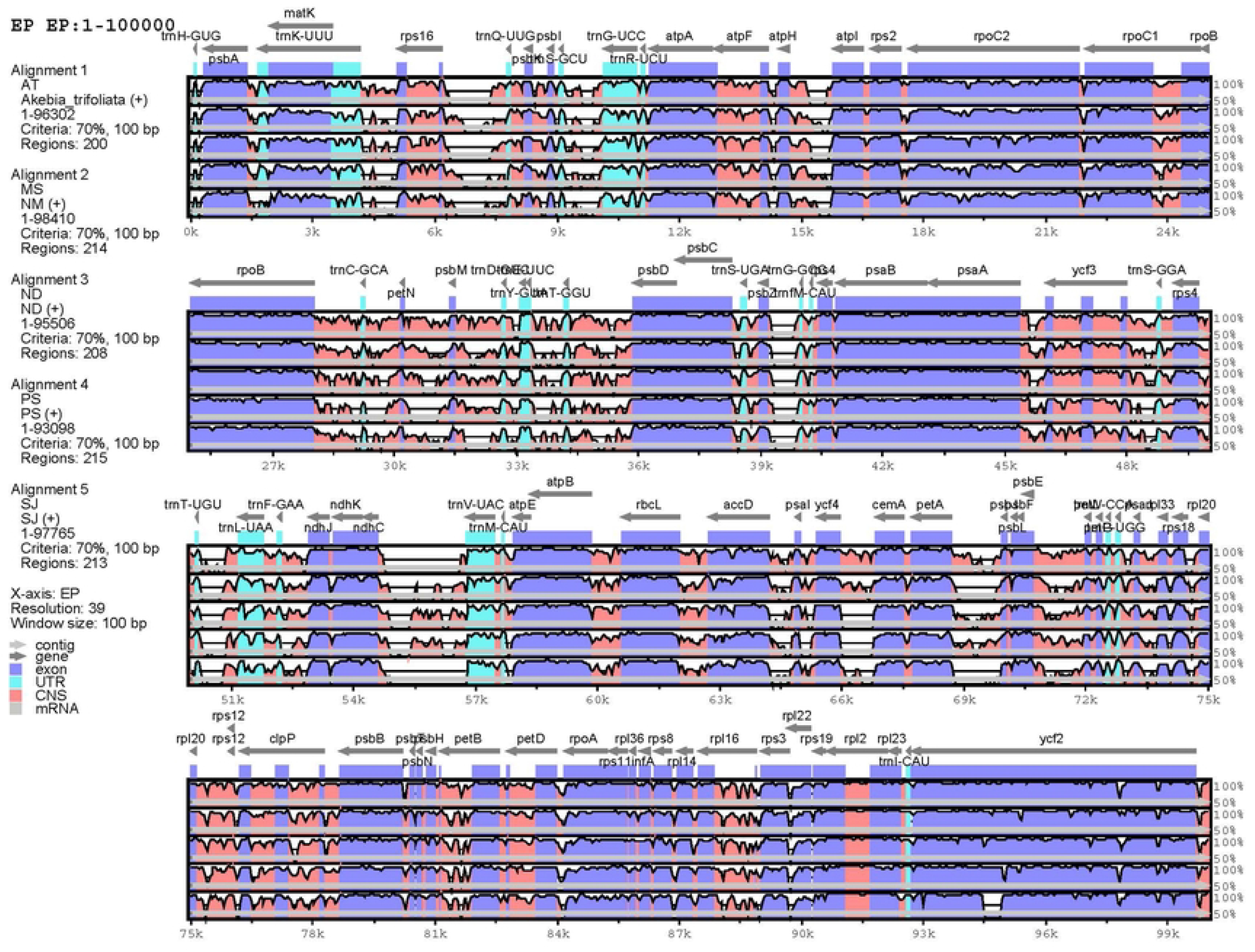
Comparison of the six chloroplast genomes using mVISTA. EP represent *Euptelea pleiospperma*, MS represent *Megaleranthis saniculifolia*, ND represent *Nandina domestica*, PS represent *Papaver somniferum*, SJ represent *Stephania japonica*. Gray arrows and thick black lines above the alignment indicate gene orientation. Purple bars represent exons, blue bars represent untranslated regions (UTRs), pink bars represent conserved non-coding sequences (CNS), and gray bars represent mRNA. The y-axis represents the percentage identity (shown: 50-70%).

74 protein-coding genes from 41 angiosperm species that belonging to 15 commelinid-clade orders were used to reconstructed the phylogenetic tree based on the maximum likelihood method. The evolutionary relationship among angiosperms is presented with high bootstrap supports (Figure 4). It shown that *A. trifoliata* is closely related to *S. japonica*, which then formed a cluster with *N. dormestica, M. saniculifolia* with 100% bootstrap supports, and these taxa belong to Ranunculales.

**Figure 4.**
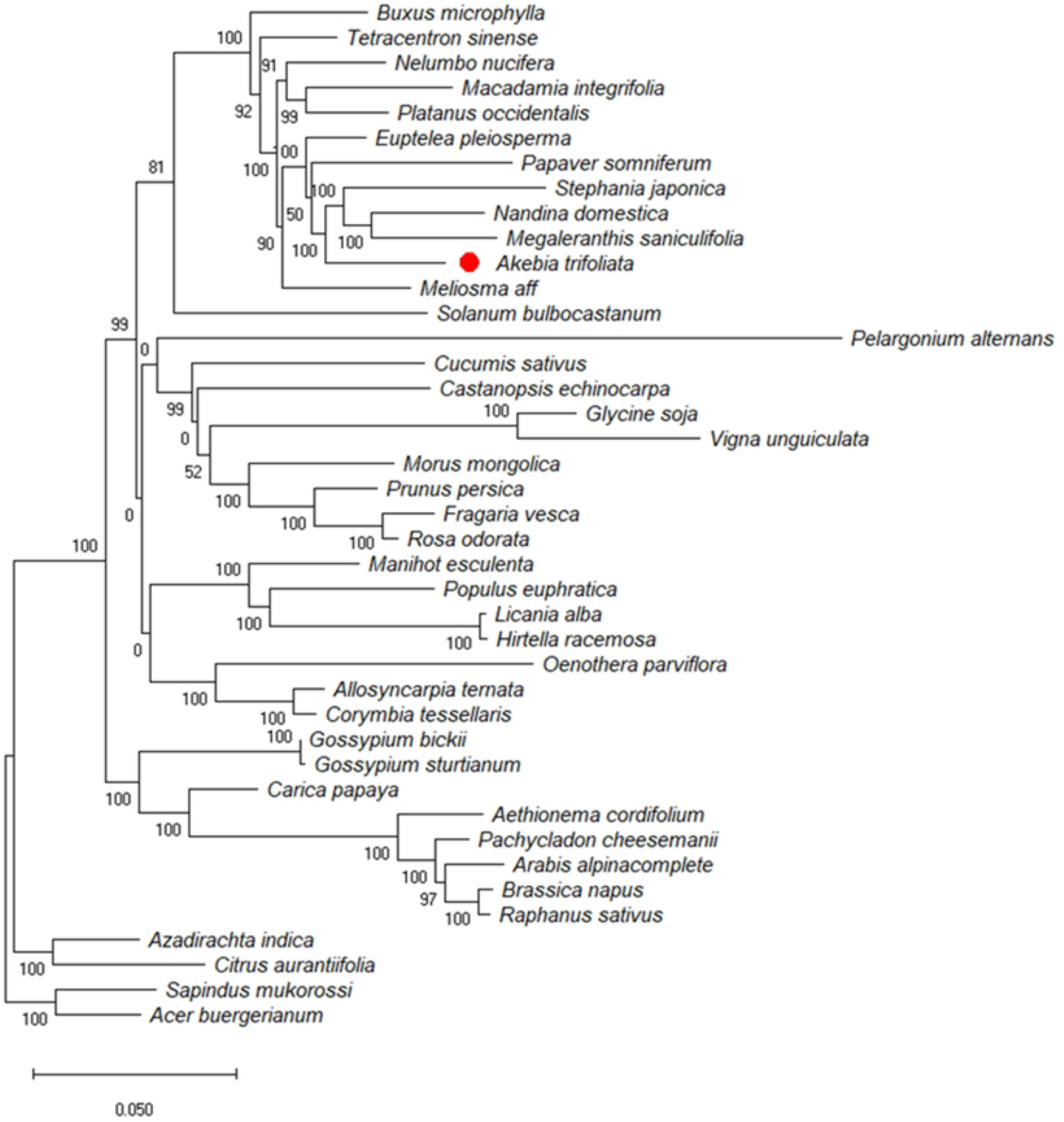
Maximum likelihood (ML) phylogenetic tree reconstruction including 41 species based on concatenated sequences from all chloroplast genomes. The position of *A. trifoliata* is indicated by red bar. *S. mukorossi* and *A. buergerianum* from Sapindales were used as outgroups.

## Discussion

With the development of NGS, chloroplast genome sequences can be obtained efficiently and economically. In the present study, we obtained the complete sequence of the *A. trifoliata* chloroplast genome, which was fully characterized and compared to the chloroplast genomes of species from different orders. *The A. trifoliata* chloroplast genome includes 130 unique genes encoding 80 proteins, 8 rRNAs, 37 tRNAs and two 2 pseudogenes. The obtained chloroplast genome of *A. trifoliata* has a typical quadripartite structure, and its gene content, gene order and GC content are similar to those of most other species from different orders.

Simple sequence repeats (SSRs) are significant repetitive elements of the entire genome and play important roles in genome recombination and rearrangement. The SSRs in chloroplast genomes are usually distributed in intergenic regions. In the SSR analysis, 108 SSR loci were found, and most SSRs were located in the intergenic region. The SSRs identified in the chloroplast genome of Akebia trifoliata can be used to analyze polymorphisms at the intraspecific level. They can also be used to develop lineage-specific markers for future evolutionary and genetic diversity studies. The mVISTA results showed that the sequence of the chloroplast genome is highly conserved among the Ranunculales clade. In addition, it showed that the sequence and content of IR regions are more conserved than are those of the LSC and SSC regions among the studied species, possibly because of the rRNA in IR regions.

In this study, the phylogenetic position of *A. trifoliata* in the Malpighiales was inferred by analyzing the complete chloroplast genome. The results suggest that *A. trifoliata* is most closely related to *Stephania japonica*, which then formed a cluster with *Nandina dormestica, Megaleranthis saniculifolia* to form subgroup of Ranunculales. Our study on the Akebia trifoliata chloroplast genome provides information on mangrove plant species in coastal intertidal zones. Moreover, the chloroplast genomic data provided in this study will be valuable for future phylogenetic studies and other studies of mangrove species.

## Conclusions

We successfully assembled, annotated and analyzed the complete chloroplast sequence of *A. trifoliata*, a Ranales clade species. The chloroplast genome was found to be conserved among several species. We identified 168 SSR loci and 23 long repeats in the chloroplast, which can be used for the development of lineage-specific markers. The LSC/IRB/SSC/IRA boundary regions of the chloroplast genome were compared among six species, and the results revealed that all these cp genomes appeared to be structurally similar with a typical quadripartite structure of two IRs separated by an LSC and an SSC. The phylogenetic analyses showed that *A. trifoliata* is most closely related to *Stephania japonica* among the order of Ranunculales. The molecular data in this study represent a valuable resource for the study of evolution in mangrove species.

## Acknowledgements

This work was supported by Research and Demonstration on Key Technologies of Ecological Planting Authentic Chinese Medicinal Materials in Ankang (19JC001) and The Base Construction and key Technologies and Industrialization Demonstration on Suitable Chinese Herbal Medicine (2020QFY11-01). We thank Dr. WANG Tingzhang, in Zhejiang Institute of Microbiology, for his kind help in high throughput sequencing and genome sequence assembly.

## References

1. Li L, Xu Q, Yao X. Microsatellite analysis reveals the resilience of genetic diversity within extant populations of three Akebia species to chronic forest fragmentation in China. Plant Ecology. 2019;220(1):69–81.

2. Shuaiyu Z, Xiaohong Y, Caihong Z, Tingting Z, Hongwen H. Genetic analysis of fruit traits and selection of superior clonal lines in Akebia trifoliate (Lardizabalaceae). Euphytica. 2018;214(7):111.

3. Li L, Yao X, Zhong C, Chen X, Huang H. Akebia: A Potential New Fruit Crop in China. Hortence A Publication of the American Society for Horticultural ence. 2010;45(1):4–10.

4. Zhongyan W, Caihong Z, Fanwen B, Difei P, Juncai P, Feirong Y. Akebia - a valuable wild fruit under domestication. Agricultural Science & Technology. 2005;6(2):12–7.

5. Wang J, Xu QL, Zheng MF, Ren H, Lei T, Wu P, et al. Bioactive 30-noroleanane triterpenes from the pericarps of Akebia trifoliata. Molecules (Basel, Switzerland). 2014;19(4):4301–12. Epub 2014/04/10. doi: 10.3390/molecules19044301. PubMed PMID: 24714192; PubMed Central PMCID: PMCPMC6271860.

6. Li L, Xu-Zhong C, Xiao-Hong Y, Hua T, Hong-Wen H. Geographic Distribution and Resource Status of Three Important Akebia Species. Plant Science Journal 2010. p. 497–506.

7. Gao H, Wang Z. Triterpenoid saponins and phenylethanoid glycosides from stem of Akebia trifoliata var. australis. Phytochemistry (Amsterdam) 2006. p. 2697–705.

8. Guan S-H, Xia J-M, Lu Z-Q, Chen G-T, Jiang B-H, Liu X, et al. Structure elucidation and NMR spectral assignments of three new lignan glycosides from Akebia trifoliata. Magn Reson Chem. 2008;46(2):186–90. Epub 2007/12/21. doi: 10.1002/mrc.2152. PubMed PMID: 18095263.

9. Iwanaga, Warashina S, Miyase T, Toshio. Triterpene saponins from the pericarps of Akebia trifoliata. Chemical and Pharmaceutical Bulletin 2012. p. Chemical and Pharmaceutical Bulletin.

10. Mimaki Y. Triterpenes and Triterpene Saponins from the Stems of Akebia trifoliata. Chemical and Pharmaceutical Bulletin. 2003;8(51):960–5.

11. Shu-guang G, Wen-bo Y, Shu-hong G. The phenolic acid and phenoglycoside of Akebia trifoliata. Lishizhen Med Mat Med Res 2010. p. 905–6.

12. Neuhaus HE, Emes MJ. Nonphotosynthetic metabolism in plastids. Annual review of plant physiology and plant molecular biology. 2000;51(1):111–40. doi: 10.1146/annurev.arplant.51.1.111. PubMed PMID: 15012188.

13. Chumley TW, Palmer JD, Mower JP, Fourcade HM, Calie PJ, Boore JL, et al. The Complete Chloroplast Genome Sequence of Pelargonium × hortorum: Organization and Evolution of the Largest and Most Highly Rearranged Chloroplast Genome of Land Plants. Molecular biology and evolution. 2006;23(11):2175–90. doi: 10.1093/molbev/msl089.

14. Lin C-P, Huang J-P, Wu C-S, Hsu C-Y, Chaw S-M. Comparative Chloroplast Genomics Reveals the Evolution of Pinaceae Genera and Subfamilies. Genome biology and evolution. 2010;2:504–17. doi: 10.1093/gbe/evq036.

15. Petit RJ, Duminil J, Fineschi S, Hampe A, Salvini D, Vendramin GG. Comparative organization of chloroplast, mitochondrial and nuclear diversity in plant populations. Molecular ecology. 2005;14(3):689–701. Epub 2005/02/23. doi: 10.1111/j.1365-294X.2004.02410.x. PubMed PMID: 15723661.

16. Hirao T, Watanabe A, Kurita M, Kondo T, Takata K. Complete nucleotide sequence of the Cryptomeria japonicaD. Don. chloroplast genome and comparative chloroplast genomics: diversified genomic structure of coniferous species. BMC plant biology. 2008;8(1):70. doi: 10.1186/1471-2229-8-70.

17. Yi X, Gao L, Wang B, Su YJ, Wang T. The complete chloroplast genome sequence of Cephalotaxus oliveri (Cephalotaxaceae): evolutionary comparison of cephalotaxus chloroplast DNAs and insights into the loss of inverted repeat copies in gymnosperms. Genome biology and evolution. 2013;5(4):688–98. Epub 2013/03/30. doi: 10.1093/gbe/evt042. PubMed PMID: 23538991; PubMed Central PMCID: PMCPMC3641632.

18. Chang CC, Lin HC, Lin IP, Chow TY, Chen HH, Chen WH, et al. The chloroplast genome of Phalaenopsis aphrodite (Orchidaceae): comparative analysis of evolutionary rate with that of grasses and its phylogenetic implications. Molecular biology and evolution. 2006;23(2):279–91. Epub 2005/10/07. doi: 10.1093/molbev/msj029. PubMed PMID: 16207935.

19. Wicke S, Schneeweiss GM, dePamphilis CW, Muller KF, Quandt D. The evolution of the plastid chromosome in land plants: gene content, gene order, gene function. Plant molecular biology. 2011;76(3-5):273–97. Epub 2011/03/23. doi: 10.1007/s11103-011-9762-4. PubMed PMID: 21424877; PubMed Central PMCID: PMCPMC3104136.

20. Shinozaki K, Ohme M, Tanaka M, Wakasugi T, Hayshida N, Matsubayasha T, et al. The complete nucleotide sequence of the tobacco chloroplast genome. Plant Molecular Biology Reporter. 1986;4(3):111–48. doi: 10.1007/BF02669253.

21. Ohyama K, Fukuzawa H, Kohchi T, Shirai H, Sano T, Sano S, et al. Chloroplast gene organization deduced from complete sequence of liverwort Marchantia polymorpha chloroplast DNA. Nature. 1986;322(6079):572–4. doi: 10.1038/322572a0.

22. Thiel T, Michalek W, Varshney RK, Graner A. Exploiting EST databases for the development and characterization of gene-derived SSR-markers in barley (Hordeum vulgare L.). TAG Theoretical and applied genetics Theoretische und angewandte Genetik. 2003;106(3):411–22. Epub 2003/02/18. doi: 10.1007/s00122-002-1031-0. PubMed PMID: 12589540.

23. Kurtz S, Choudhuri JV, Ohlebusch E, Schleiermacher C, Stoye J, Giegerich R. REPuter: the manifold applications of repeat analysis on a genomic scale. Nucleic acids research. 2001;29(22):4633–42. Epub 2001/11/20. doi: 10.1093/nar/29.22.4633. PubMed PMID: 11713313; PubMed Central PMCID: PMCPMC92531.

24. Frazer KA, Pachter L, Poliakov A, Rubin EM, Dubchak I. VISTA: computational tools for comparative genomics. Nucleic acids research. 2004;32(Web Server issue):W273–9. Epub 2004/06/25. doi: 10.1093/nar/gkh458. PubMed PMID: 15215394; PubMed Central PMCID: PMCPMC441596.

